# Ubiquitous overexpression of CXCL12 confers radiation protection and enhances mobilization of hematopoietic stem and progenitor cells

**DOI:** 10.1101/2020.01.08.899427

**Authors:** Smrithi Rajendiran, Stephanie Smith-Berdan, Leo Kunz, Maurizio Risolino, Licia Selleri, Timm Schroeder, E Camilla Forsberg

**Affiliations:** Institute for the Biology of Stem Cells, Department of Biomolecular Engineering, University of California Santa Cruz, Santa Cruz, CA 95064, USA; Department of Biosystems Science and Engineering, Eidgenössische Technische Hochschule Zürich, 4058 Basel, Switzerland; Program in Craniofacial Biology, Institute of Human Genetics, Eli and Edyth Broad Center of Regeneration Medicine & Stem Cell Research, Departments of Orofacial Sciences and Anatomy, University of California, San Francisco, 513 Parnassus Avenue, HSW 710, San Francisco, CA 94143, USA

**Author notes:** Corresponding author: Director, Institute for the Biology of Stem Cells, Professor of Biomolecular Engineering, University of California Santa Cruz, Mail stop SOE2, 1156 High Street, Santa Cruz, CA 95064, USA.

**Keywords:** Hematopoietic stem cells, CXCL12 transgenic, hematopoiesis, radiation protection, mobilization.

## Abstract

C-X-C Motif Chemokine Ligand 12 (CXCL12; aka SDF1α) is a major regulator of a number of cellular systems, including hematopoiesis where it influences hematopoietic cell trafficking, proliferation, and survival during homeostasis and upon stress and disease. A variety of constitutive, temporal, ubiquitous and cell-specific loss-of-function models have documented the functional consequences on hematopoiesis upon deletion of *Cxcl12*. Here, in contrast to loss-of-function experiments, we implemented a gain-of-function approach by generating a dox-inducible transgenic mouse model that enables spatial and temporal overexpression of *Cxcl12*. We demonstrated that ubiquitous CXCL12 overexpression led to an increase in multipotent progenitors in the bone marrow and spleen. The CXCL12+ mice displayed reduced reconstitution potential as either donors or recipients in transplantation experiments. Additionally, we discovered that *Cxcl12* overexpression improved hematopoietic stem and progenitor cell mobilization into the blood, and conferred radioprotection by promoting quiescence. Thus, this new CXCL12+ mouse model provided new insights on major facets of hematopoiesis and serves as a versatile resource for studying CXCL12 function in a variety of contexts.

## INTRODUCTION

Cytokines and chemokines play essential roles in a number of important biological processes (Ahmed et al., 2014; Chen et al., 2018; Doring et al., 2014; Edderkaoui, 2017; Janssens et al., 2018; Ridiandries et al., 2018; Sokol and Luster, 2015; Turner et al., 2014), including hematopoiesis (Wright et al., 2002; Youn et al., 2000) and cancer metastasis (Agarwal et al., 2019; Amedei et al., 2013; Bridge et al., 2018; Chow and Luster, 2014; King et al., 2017; Marcuzzi et al., 2018; Nagarsheth et al., 2017). Because of its clearly demonstrated essential roles for hematopoiesis, CXCL12 is one of the most intensely studied CXC subfamily chemokine factor in the bone marrow (BM) niche (Abe-Suzuki et al., 2014; Aurrand-Lions and Mancini, 2018; Broxmeyer and Kim, 1999; Lewellis and Knaut, 2012; Sugiyama et al., 2006). During development, CXCL12 is essential for efficient migration of hematopoietic stem cells (HSCs) from the fetal liver, the primary hematopoietic organ during late-gestation mammalian development, to the BM, the main site of post-birth hematopoiesis (Ara et al., 2003; Christensen et al., 2004). Loss of function studies in the mouse by deletion of *Cxcl12*, or its primary receptor *Cxcr4*, results in lethality which could at least in part be contributed by defects in hematopoiesis, in addition to other aberrations in cardiac septum formation, neuronal cell migration and vasculogenesis (Ma et al., 1998; Nagasawa et al., 1996; Ratajczak et al., 2006; Tachibana et al., 1998). Perturbation of *Cxcl12* has also been associated extensively with B-cell lymphopoiesis (Nagasawa et al., 1996) and various cancers including pancreatic, breast, brain, thyroid, prostate, skin (Ahirwar et al., 2018; Domanska et al., 2013; Goffart et al., 2017; Guo et al., 2016; Liu et al., 2014; Meng et al., 2018; Patrussi and Baldari, 2011; Singh et al., 2004; Teicher and Fricker, 2010; Wald et al., 2013; Werner et al., 2017; Zboralski et al., 2017; Zhang et al., 2018). Disruption of the CXCL12/CXCR4 axis promotes mobilization of hematopoietic stem and progenitor cells (HSPCs) from the BM to the blood, facilitating the process of cell collection for transplantation (Broxmeyer et al., 2005, 2007; Dar et al., 2011; Karpova et al., 2017; Levesque et al., 2003; Smith-Berdan et al., 2011, 2015, 2019; Sugiyama et al., 2006).

Ding et al. and Greenbaum et al. have shown that cell type-specific deletion of CXCL12 in the BM results has selective effects on different types of HSPCs, suggesting that proximity to *Cxcl12*-expressing cells influences HSPC function (Ding and Morrison, 2013; Greenbaum et al., 2013; Ugarte and Forsberg, 2013). Altering the CXCL12 levels by transgenic overexpression or exogenous addition of CXCL12 has been shown to not only increase cell survival, specifically of the myeloid lineage, but also to enhance vascularization (Broxmeyer et al., 2003a). CXCL12 expression is increased upon irradiation in mice and zebrafish (Chang et al., 2009, 2018; Glass et al., 2011; Smith-Berdan et al., 2012). While loss-of-function and *in vitro* studies have shown that CXCL12 promotes quiescence of HSCs, it has also been demonstrated that CXCL12 improves the homing of HSCs to the BM in a transplantation setting (Asada et al., 2017; Nie et al., 2008; Perlin et al., 2017; Wright et al., 2002).

The aim of this study is to test the effects of elevated *Cxcl12* levels in vivo. To address this, we generated a transgenic mouse model that enables temporal and cell type-specific overexpression of CXCL12. Here, we use this model in adult hematopoiesis, without affecting the key role that CXCL12 plays in the establishment of the hematopoietic niche during gestation and initial postnatal phases.

## RESULTS

### Generation of *Cxcl12* transgenic mice

To study the role of CXCL12 on hematopoiesis, we generated a doxycycline (dox) inducible transgenic mouse by pronuclear injection of the vector shown in Figure S1A into zygotes. The construct was designed with tetracycline (tet) responsive elements (TRE) preceding the *Cxcl12* coding sequence, followed by a T2A cleavage site and a tdTomato (Tom) reporter (Figure 1A). SignalP software predicted a secreted CXCL12 protein (Figure S1B). Progeny of *Cxcl12* transgenic mice were crossed to *Rosa rtTA* mice (Hochedlinger et al., 2005). In this tet-On system, doxycycline (dox) exposure led to widespread expression of secreted CXCL12 and cellular membrane bound Tom labeling (Figure 1A). Dox-response kinetics studies showed the induction of Tom in peripheral white blood cells when mice bearing both the *Cxcl12* transgene and *Rosa rtTA* were given dox (mouse #1). The fraction of Tom+ cells decreased a few days after dox withdrawal. Additionally, no Tom+ cells were observed in mice bearing only the *Cxcl12* transgene (mouse #2) or only *rtTA* (mouse #3) (Figure S1C and S1D). Henceforth, we used the mice that have at least 10-15% of Tom positive peripheral blood cells upon dox treatment as *Cxcl12*-overexpressing mice (TgCXCL12_RosaOE_ or CXCL12+) and the transgene negative (*Rosa rtTA*, C57BL6, or mT/mG) mice that have been subjected to dox treatment at the same concentration and for the same duration as control mice (Control). To test whether Tom fluorescence is an accurate indicator of *Cxcl12* overexpression, we performed two-step qRT-PCR for the mRNA levels of *Cxcl12* in the CXCL12+ mice after dox treatment. While the qRT-PCR showed that the levels of endogenous (endo) *Cxcl12* and of its receptors, *Cxcr4* and *Cxcr7*, were not altered, the *Cxcl12* CDS (coding) mRNA was significantly increased in cells from different tissues (Figure 1B), confirming that Tom expression can be used as a convenient proxy for *Cxcl12* overexpression. Additionally, using proximity ligation assay (PLA)-based single molecule imaging, we also observed significantly elevated CXCL12 signal in the BM of CXCL12+ mice compared to controls (Figure 1C and S1E). This demonstrates increased levels of CXCL12 protein, in addition to mRNA (Figure 1B and S1F). Secretion of functional CXCL12 was tested by transwell migration assays (Smith-Berdan et al., 2011), where we observed a significant increase in the migration of HSPCs towards the BM supernatant from CXCL12+ mice compared to controls (Figure 1D). Together, these results demonstrate that the CXCL12+ mice over-produce functional CXCL12 protein in response to dox. Notably, these mice do not exhibit gross abnormalities, are viable, and fertile.

**Figure 1:**
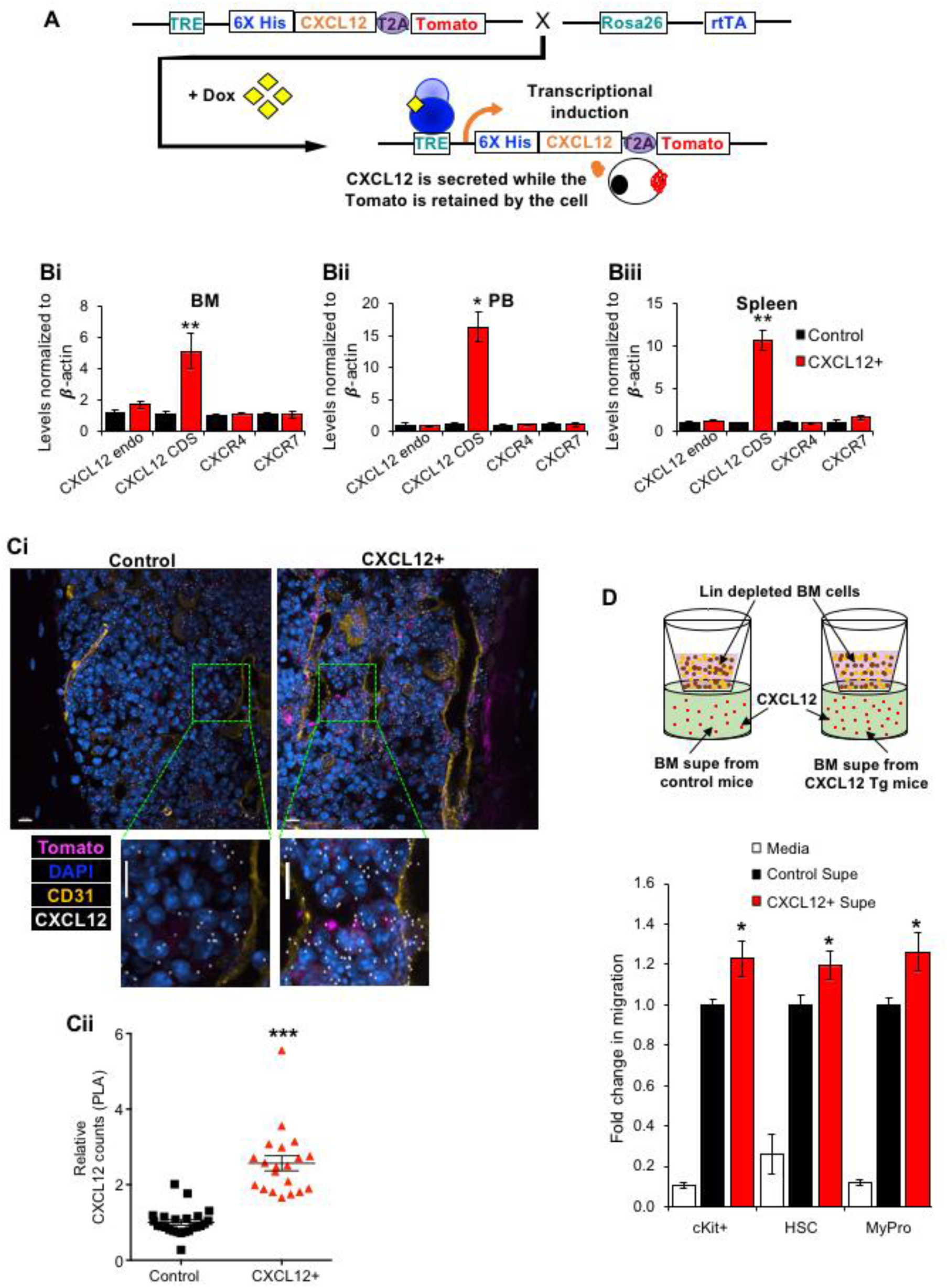
TgCXCL12_RosaOE_ mice express higher levels of CXCL12 upon dox treatment. **(A):** Strategy for generation of the *Cxcl12* transgenic mouse line, which, upon addition of doxycycline, broadly expresses CXCL12 (secreted) and tomato (retained in the cellular membrane). **(B):** RT-qPCR analysis of tissues from *Cxcl12* transgenic mice (CXCL12+) compared to controls (Control) for endogenous *Cxcl12* only (endo), *Cxcl12* endogenous and transgenic (CDS), *Cxcr4* and *Cxcr7* transcript levels. 3 independent experiments for BM (**Bi**; Control = 8 mice, CXCL12+ = 12 mice), 1 experiment each for PB (**Bii**) and Spleen (**Biii**) (Control = 2 mice, CXCL12+ = 3 mice); All the mice were maintained on Doxycycline for at least 3 days prior to tissue extraction. **(C):** Representative BM sections by PLA assay confocal imaging (**i**) depicting increased expression (scale bar depicts 10μm) (**ii**) of CXCL12 in CXCL12+ mice compared to control mice. Data from 4 sections per mouse with 6 control and 5 CXCL12+ mice. **(D):** Transwell migration assay shows increased migration of lineage-depleted BM cells towards BM supernatant from CXCL12+ mice (CXCL12+ Supe) compared to BM supernatant from controls (Control Supe). Bar graph depicts fold change in migration of cKit+, HSC and Myeloid Progenitor (MyPro) cells towards the different supernatants. Data represented from 8, 21, and 26 mice for media, control and CXCL12+ supe groups respectively, over 3 independent experiments. P-values: * ≤ 0.05; ** ≤ 0.01; *** ≤ 0.001

### TgCXCL12_RosaOE_ mice have increased numbers of multipotent progenitors

To determine the impact of CXCL12 overexpression on steady-state hematopoiesis, we quantified the total number of cells in the main hematopoietic compartments. The ubiquitous overexpression of CXCL12 led to a significant increase in multipotent progenitors (MPPs) in both the BM (Figure 2A, S2A) and the spleen (Figure 2B), which is consequently reflected as an increase in the KLS compartment (Figure S2B-C). The increase in the MPP numbers is potentially a result of either a transient increased production or a reduced differentiation rate of MPPs, or a combination of the two, without compromising HSC numbers; but is not due to the redistribution of the cells from the other major hematopoietic compartments since there is no significant deficit in any of the three tissues. No other major differences were detected in the hematopoietic (Figure 2A-C, S2B-D) or the stromal compartments (Figure S2E-F).

**Figure 2:**
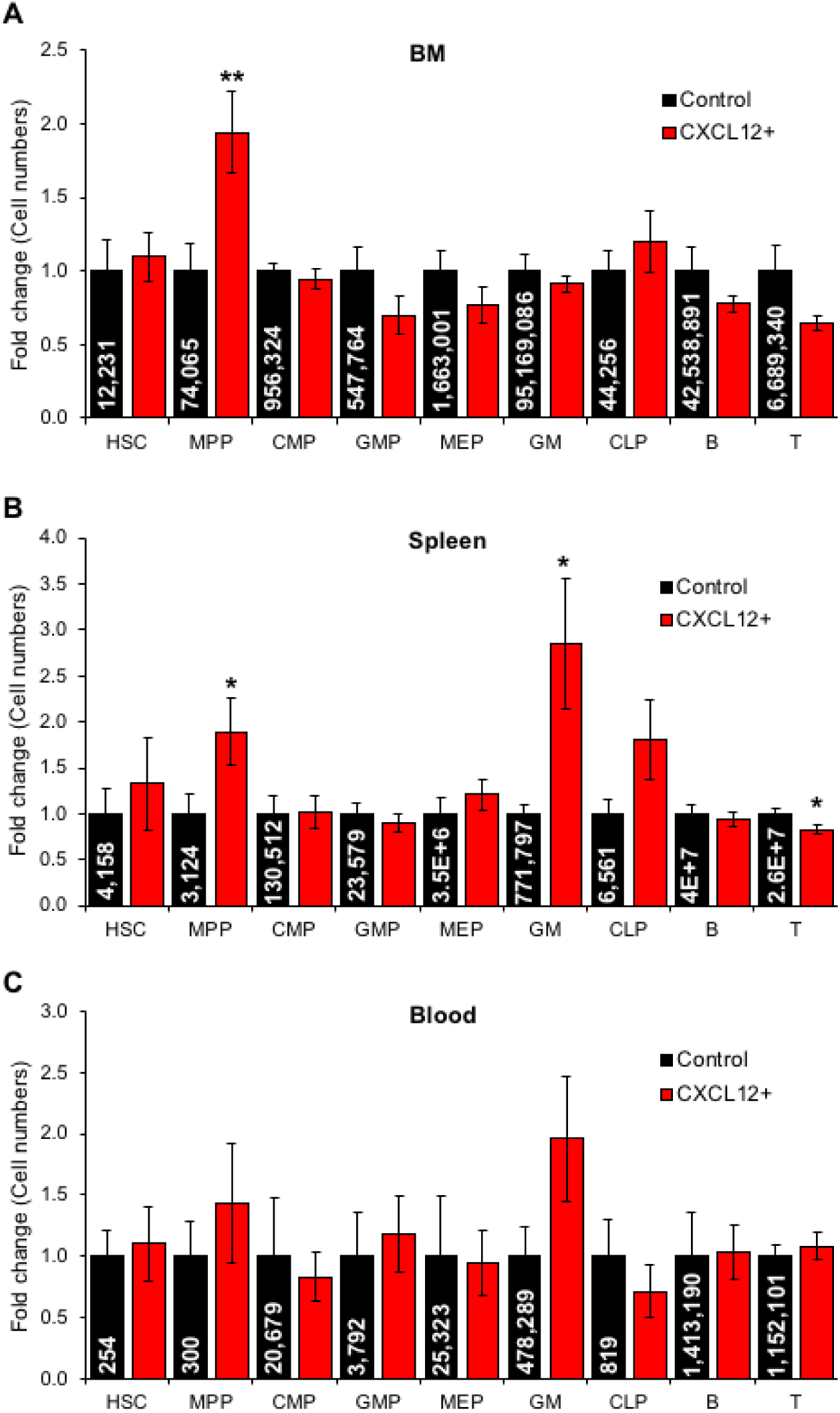
Ubiquitous overexpression of *Cxcl12* increases the numbers of MPP in the BM and spleen. **(A-C):** HSPCs and Mature cells in the BM (**A**), Spleen (**B**) and Blood (**C**) of control and CXCL12+ mice. Mice were given Doxycycline for at least 7 days prior to perfusion and tissue extraction to obtain the different cell types. Data represented from at least 15 mice in each group over 3 independent experiments. Numbers in the bars represent absolute detected cell counts in the mouse BM (assuming BM from both the femurs and tibia account for 25% of the total BM); entire spleen and blood. P-values: * ≤ 0.05; ** ≤ 0.01

### HSCs exposed to higher CXCL12 concentrations prior to transplantation show reduced reconstitution potential

In order to determine if the HSCs in CXCL12+ mice have differential long-term multi-lineage reconstitution (LTMR) potential, we transplanted HSCs isolated from CXCL12+ mice or control mice into UBC-GFP mice (Figure 3A). Interestingly, engraftment from HSCs that had been exposed to high levels of CXCL12 pre-transplantation was lower than that obtained with HSCs from control mice, measured both as donor chimerism (Figure 3Bi) and as absolute cell numbers (Figure 3Bii). Thus, increased levels of CXCL12 (Figure 3) decrease the LTMR potential of HSCs. Of note, it was previously described that reduced CXCL12 levels also reduce the LTMR potential of HSCs (Ding and Morrison, 2013).

**Figure 3:**
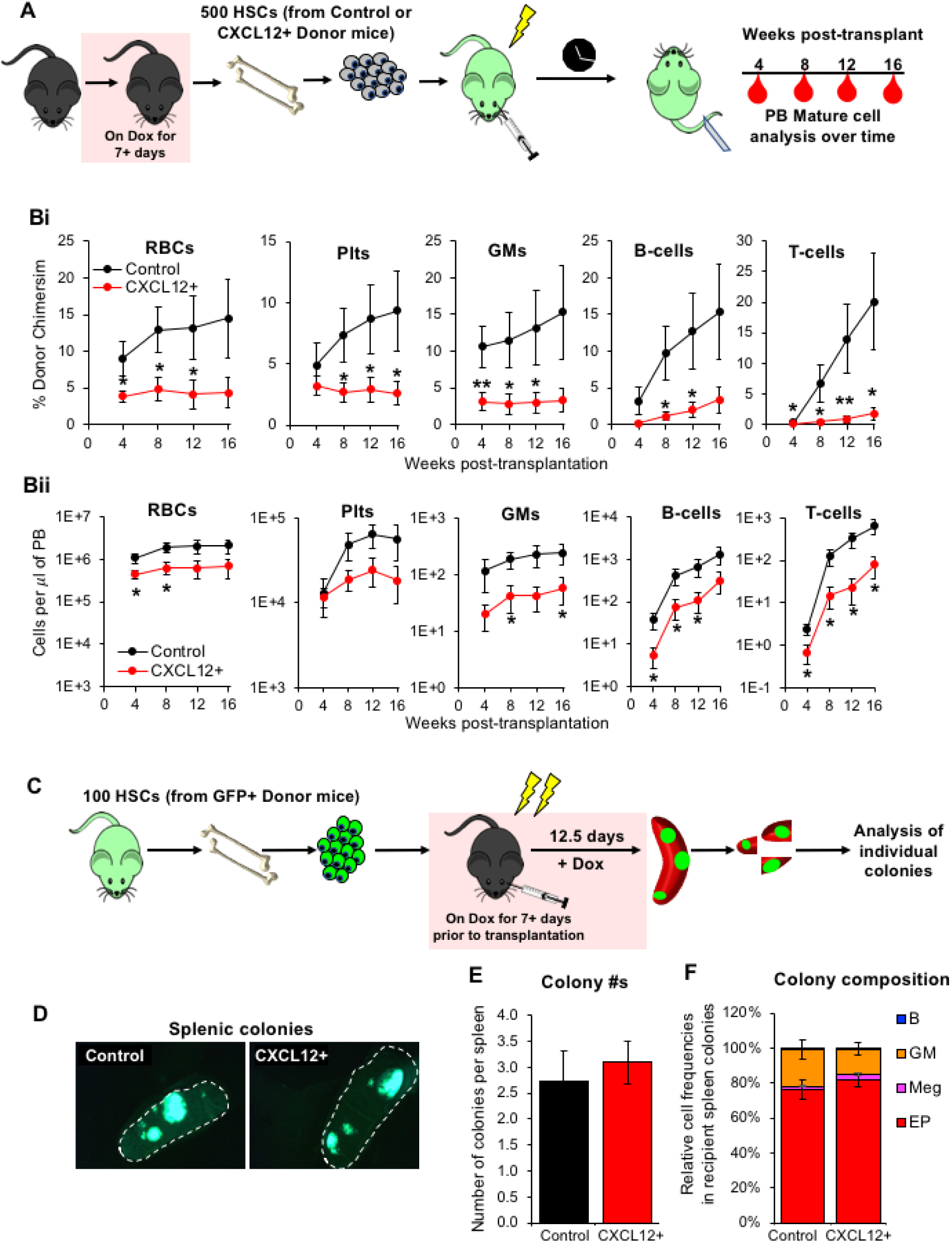
Repopulation by HSCs from CXCL12+ mice in wild-type recipients is lower than that obtained using HSCs from control mice. **(A):** Schematic showing transplantation of 500 HSCs from control or CXCL12+ mice into sub-lethally (525 rads) irradiated UBC-GFP WT recipient mice to obtain long term multilineage reconstitution in the PB over time. **(B):** Peripheral blood reconstitution assessed by donor chimerism (**i**) and absolute numbers (**ii**) for each of the mature cell types; i.e, RBCs (Ter119+ CD61- Mac1- Gr1- B220- CD3-), Platelets (Ter119-CD61+ Mac1- Gr1- B220- CD3-), GMs (Ter119- CD61- Mac1+ Gr1+ B220- CD3-), B-cells (Ter119- CD61- Mac1- Gr1- B220+ CD3-) and T-cells (Ter119- CD61- Mac1- Gr1- B220- CD3+). Data from 2 independent experiments with 11 control and 15 CXCL12+ mice. **(C):** Study design for transplantation of 100 WT GFP+ HSCs into lethally irradiated (1050 rads) control or CXCL12+ mice. **(D):** Representative images of GFP+ colonies in the spleens of mice 12.5 days post transplantation. **(E):** Relative number of colonies per spleen in control and CXCL12+ mice. **(F):** Distribution of cell types within the colonies. Data from 2 independent experiments with 11 mice per group. Red boxes indicate mice on Dox (1mg/ml) with 5% Sucrose. P-values: * ≤ 0.05; ** ≤ 0.01

### Increased CXCL12 in the recipient does not differentially regulate homing of HSPCs

CXCL12 produced by CXCL12-abundant reticulocytes (CAR cells) in the BM has been associated with the retention of HSCs to the BM (Asada et al., 2017; Sugiyama et al., 2006). In addition, loss of CXCL12 significantly reduces HSC engraftment in the BM (Ding and Morrison, 2013). Recently, we demonstrated by quantitative imaging that *Cxcl12* is enriched at putative HSC niches, but does not form protein gradients towards them (Kunz and Schroeder, 2019). To determine if disruption of the normally cell type-specific CXCL12 expression pattern is conducive to engraftment of donor cells we transplanted HSPCs into our model that provides an environment of ubiquitous *Cxcl12* overexpression. We transplanted GFP+ HSCs into CXCL12+ mice and first tested extramedullary hematopoiesis in the spleen (Figure 3C). No significant differences were observed in the splenic colonies in terms of their size (Figure 3D), their numbers (Figure 3E) or their composition (Figure 3F) between controls and CXCL12+ mice that received GFP+ HSCs.

Subsequently, we performed short-term homing assay with WT GFP+ KLS cells (Figure 4A). Here, 3 hours post-transplantation into CXCL12+ or control mice, cells from the BM, blood, spleen, lung and liver were extracted to obtain WT GFP+ donor cells. Significant differences in donor cell recovery were observed only in the liver (Figure 4Bv), whereas numbers of WT donor cells in other tissues, including the BM, were not significantly different between CXCL12+ and control recipient mice (Figure 4Bi-iv). Thus, HSPC homing is not grossly abnormal upon *Cxcl12* overexpression in any of the major hematopoietic tissues.

**Figure 4:**
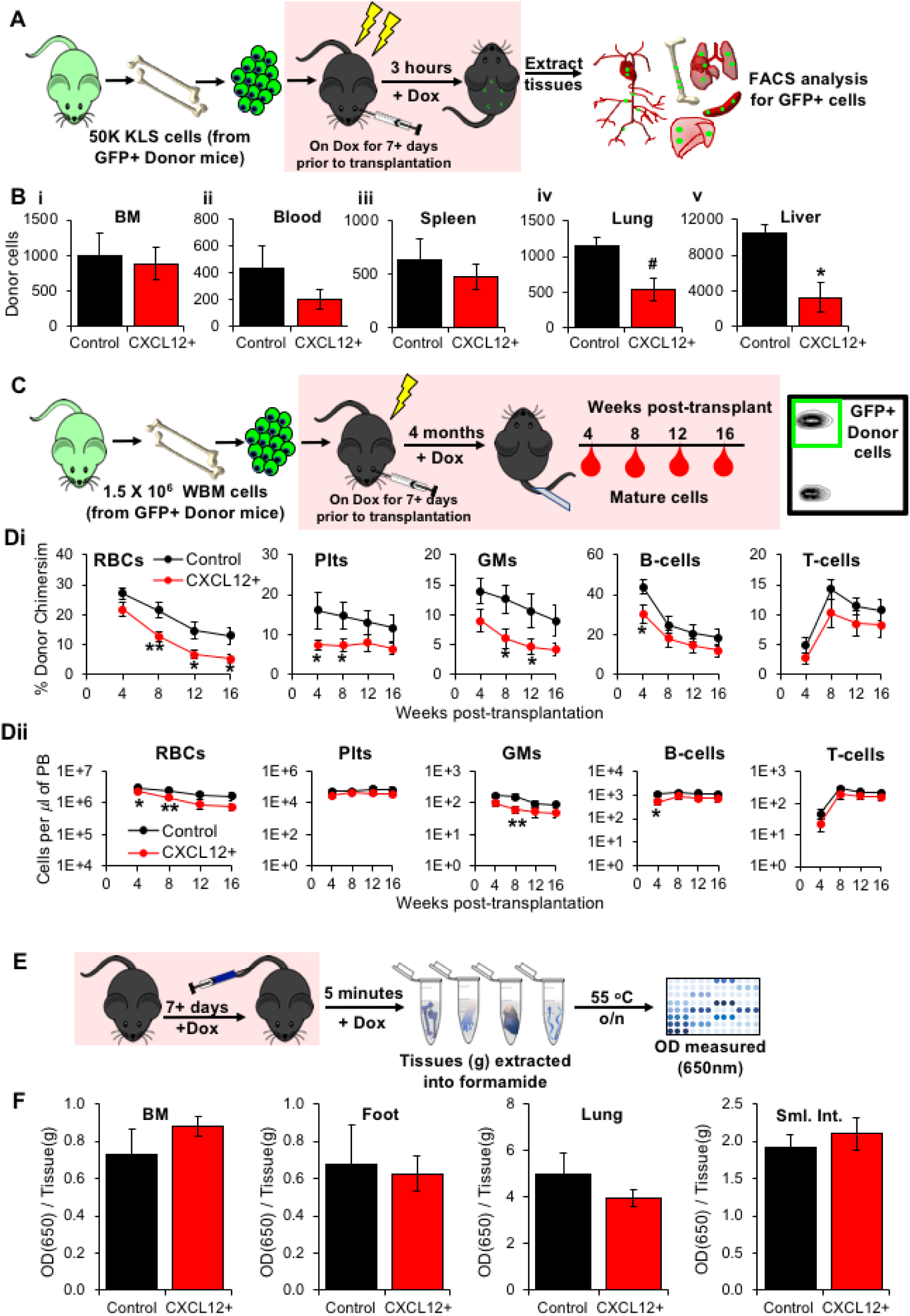
*Cxcl12* overexpression in the recipient does not affect short-term homing to the BM, but impairs long-term multilineage reconstitution in the blood. **(A)**: Schematic for short-term homing assay showing transplantation of 50K GFP+ KLS donor cells into lethally irradiated (1050 rads) control or CXCL12+ recipients and analysis of the various tissues 3 hours post-transplantation. **(B):** Donor GFP+ cells in various tissues (BM, blood, spleen, lung and liver) of control and transgenic recipients. Data from 3 independent experiments with 5 control and 11 CXCL12+ mice. **(C):** Schematic of the long-term reconstitution assay showing transplantation of GFP+ donor cells into sub-lethally irradiated (725 rads) control or CXCL12+ mice. **(D):** Peripheral blood reconstitution by donor chimerism (**i**) and absolute numbers (**ii**) for each of the mature cell types aforementioned after transplantation of 1.5 × 10_6_ WBM cells. Data from 2 independent experiments with 7 control and 11 CXCL12+ mice. **(E):** Schematic for Evans Blue (50 mg/kg) based Miles assay showing vascular permeability in the various tissues quantified as OD at 650nm per gram of tissue, 5 minutes after dye injection. **(F):** Vascular permeability is not altered upon *Cxcl12* overexpression in the BM, foot, lung or small intestine (sml. int.). Data from 4 independent experiments with 8 control and 15 CXCL12+ mice. Red boxes indicate mice on Dox (1mg/ml) with 5% Sucrose. P-value: # ≤ 0.075; * ≤ 0.05

### *Cxcl12* overexpressing niches are less conducive to the long-term engraftment of WT donor cells despite intact vasculature

Next, to determine whether ubiquitous overexpression of *Cxcl12* affected sustained long-term reconstitution, we transplanted 7.5 × 10_6_ WT GFP+ WBM cells into sub-lethally irradiated control or CXCL12+ mice (Figure S3A). No significant differences were observed between the different recipients, either in donor chimerism (Figure S3Bi) or absolute cell numbers in the blood (Figure S3Bii) over 4 months post-transplantation. However, niche space has been reported to be a limiting factor during transplantation (Bhattacharya et al., 2008; Weissman and Shizuru, 2008). Transplantation of 7.5 million WBM cells into partially cleared niches could potentially compete for the limited available space. Furthermore, elevated CXCL12 levels might not influence the engraftment of donor cells when they are introduced into the host in high numbers. Thus, to overcome these potential drawbacks and to determine the effect of *Cxcl12* overexpression in a partially cleared niche, we subsequently transplanted 1/5_th_ of the number of cells in control or CXCL12+ mice. With lower numbers of transplanted cells (1.5 × 10_6_ WBM cells per mouse) we observed a transient, but significant, decrease of engraftment in the CXCL12+ compared to control hosts (Figure 4Di-ii). Over time, the differences disappeared (Figure 4D). Similar results were obtained upon transplantation of 1000 GFP+ KLS cells (Figure S3A): initially we observed poorer multi-lineage reconstitution, but the differences between recipients disappeared over time (Figure S3C). Interestingly, while the erythro-megakaryo-myeloid lineages were typically affected at multiple timepoints, B-cell reconstitution was significantly reduced only at 4 weeks after transplantation, and T-cells were not altered (Figure 4Di-ii and S3Ci-ii). To test if the above described lineage-specific reduction is a direct consequence of *Cxcl12* overexpression that affects myeloid progenitors (MyPros) more strongly, we transplanted MyPros isolated from untreated WT mice into CXCL12+ or control mice. No differences in the reconstitution potential were revealed between the two hosts (Figure S3D), indicating that the lower engraftment observed is not due to inefficient performance of the MyPros in a *Cxcl12* overexpressing environment. We and others have shown that vascular integrity is an important regulator of HSPC location and engraftment (Ding et al., 2012; Perlin et al., 2017; Poulos et al., 2017; Smith-Berdan et al., 2015, 2019). To determine if *Cxcl12* overexpression causes vascular leak, we performed Miles assays by injecting Evans Blue dye in CXCL12+ or control mice (Figure 4E). We did not observe significant differences in vascular permeability of foot, BM, lung, or small intestine from CXCL12+ mice compared to controls (Figure 4F). This indicates that *Cxcl12* overexpression does not alter vascular permeability broadly, and that the reduced engraftment observed in CXCL12+ mice after transplantation is not a consequence of vascular leak.

### *Cxcl12* overexpression improves the mobilization of HSPCs to the blood

The CXCL12-CXCR4 interaction is critical to retain cells in the BM (Karpova et al., 2017; Levesque et al., 2003; Nie et al., 2008; Sugiyama et al., 2006). Disruption of this interaction using the CXCR4 inhibitor AMD3100 results in the egress of HSPCs from the BM to the blood (Bonig et al., 2009; Dar et al., 2011). To test the effect of widespread overexpression of *Cxcl12* on mobilization, we targeted the CXCR4-CXCL12 axis using AMD3100 (Figure 5A). As expected, AMD3100-treated control mice had increased numbers of mobilized KLS cells compared to non-mobilized mice (Figure 5B). In addition, we observed a significant increase of KLS cells in the blood from AMD3100-treated CXCL12+ mice compared to AMD3100-treated controls (Figure 5B). Similar trends were observed for both HSCs (Figure S4A) and MPPs (Figure S4B). Additionally, while comparing Lin-cKit+ cells (Figure S4C) and myeloid progenitors (Figure S4D), we observed an increase in the overall mobilization of these cells in CXCL12+ mice. These data show that widespread overexpression of *Cxcl12* further facilitates AMD3100-mediated mobilization of HSPCs from the BM to the blood.

**Figure 5:**
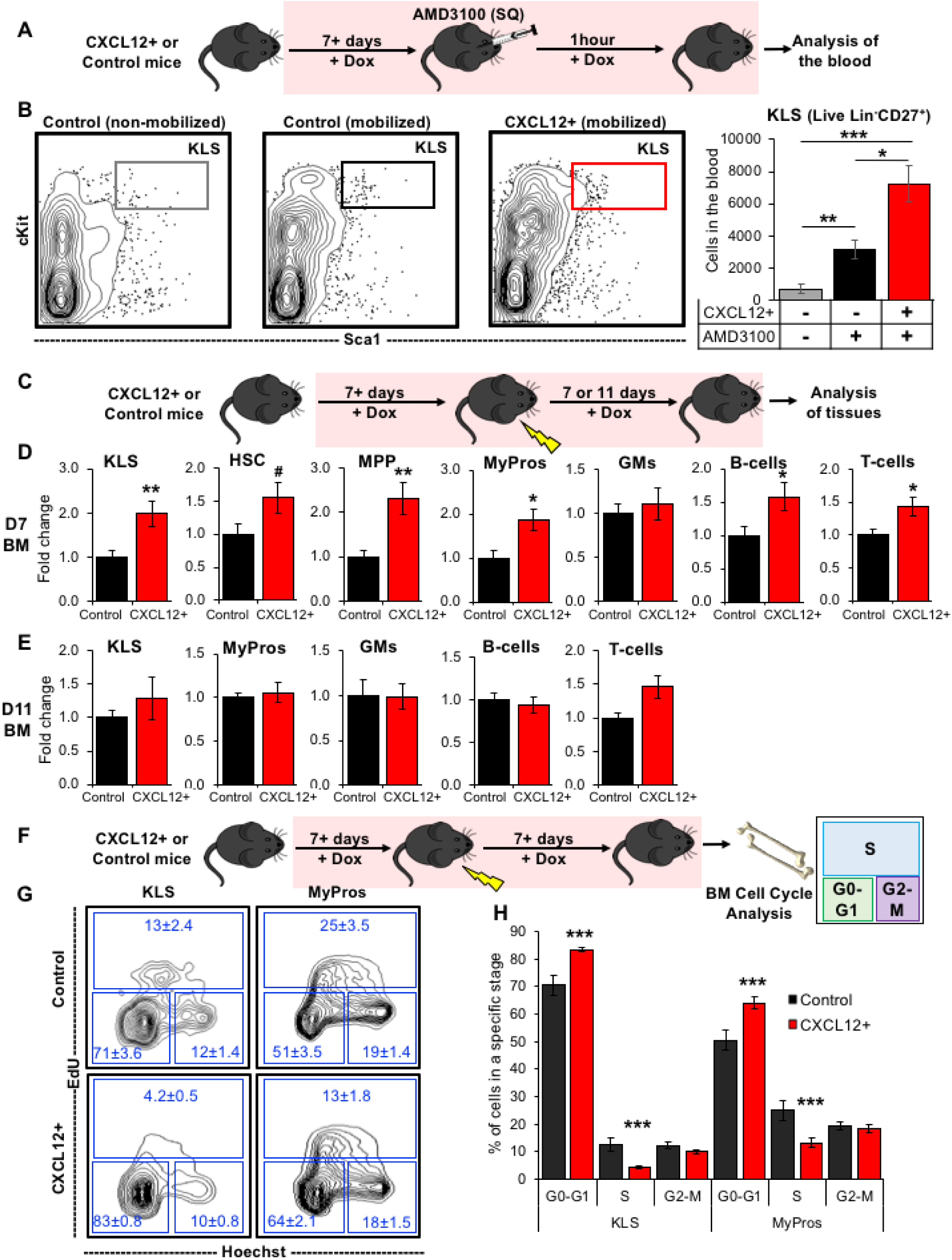
Ubiquitous overexpression of *Cxcl12* enhances AMD3100-mediated HSPC mobilization and decreases radiation-dependent HSPC ablation by promoting cell quiescence. **(A):** Control or CXCL12+ mice are injected subcutaneously with 5mg/kg AMD3100 one hour prior to analysis of HSPCs in the blood. **(B):** Representative flow plots (left) and quantification of total KLS cells (right) in the blood shows increased mobilization in CXCL12+ mice compared to controls, with and without AMD3100. Data from 4 independent experiments with 7 mice in the non-mobilized group, 7 control mice and 11 CXCL12+ mice. P-values calculated by t-test between any two groups. **(C):** Experimental design for radiation ablation. Control or CXCL12+ mice were sub-lethally irradiated (725 rads) and the different HSPCs were analyzed 7- or 11-days post irradiation. **(D):** HSPCs and mature cells in the BM, 7-days after irradiation; Fold change normalized to controls for each experiment. Data from 3 independent experiments with 12 control and 15 CXCL12+ mice. **(E):** HSPCs and mature cells in the BM, 11-days after irradiation; Fold change normalized to controls in each experiment. Data from 3 independent experiments with 16 control and 20 CXCL12+ mice. **(F):** Study design for in vivo cell cycle analysis by EdU assay. EdU is injected intraperitoneally, 30 minutes before BM extraction, into control or CXCL12+ mice that have been irradiated 7 days prior to HSPC analysis. **(G):** Representative flow plots for cell cycle analysis of KLS and MyPro cells. **(H):** Quantification of the cycling status of cKit enriched HSPCs from BM of control and CXCL12+ mice show increased cell quiescence upon *Cxcl12* overexpression. Data from 3 independent experiments with 14 control and 16 CXCL12+ mice. All tissues were analyzed after perfusion. Red boxes indicate mice on Dox (1mg/ml) with 5% Sucrose. P-values: * ≤ 0.05; ** ≤ 0.01; *** ≤ 0.001

### Overexpression of *Cxcl12* protects HSPCs against irradiation by promoting quiescence

Elevated CXCL12 has been linked to tumor progression, along with increasing resistance to radiation and chemotherapy, and to immune suppression driven by the tumor microenvironment (Duda et al., 2011; Liu et al., 2014; Zboralski et al., 2017). Our lab and others have shown that irradiation causes an elevation of CXCL12 in the BM niche (Chang et al., 2009, 2018; Smith-Berdan et al., 2012). Here, we also observed reduced post-transplantation engraftment into irradiated CXCL12+ hosts (Figure 4D and S3C). Hence, we hypothesized that overexpression of *Cxcl12* provides protection to the host cells upon irradiation stress, thereby increasing the competition between the host and donor cells post-transplantation. To test this, we irradiated CXCL12+ mice and quantified the cell numbers in BM, spleen and blood (Figure 5C). We observed an increase in HSPCs and MyPros, as well as in mature lymphoid cells (Figure 5D). The higher numbers of surviving cells in the BM of CXCL12+ mice compared to controls 7 days post-irradiation showed that CXCL12 confers protection against irradiation (Figure 5D). Similar trends were observed in the spleen and blood (Figure S4E-F), arguing against cell redistribution between tissues. By 11 days post-irradiation, cells in control and CXCL12+ mice recovered to comparable extents (Figure 5E), indicating that the D7 perturbations observed in irradiated CXCL12+ mice are either due to higher proliferation or protection from cell death as compared to control mice.

Broxmeyer et al. previously demonstrated reduced apoptosis in BM cells from *Cxcl12* transgenic mice compared to controls by decreased active Caspase-3 in the transgenic cells (Broxmeyer et al., 2003a). To test whether the radiation protection in CXCL12+ mice could be attributed to increased numbers of quiescent cells, we performed EdU-based cell cycle analysis of KLS and MyPro BM progenitor cells 7 days post-irradiation (Figure 5F). Consistent with previous data (Itkin and Lapidot, 2011; Sugiyama et al., 2006; Tzeng et al., 2011), a higher proportion of cells from CXCL12+ mice BM resided in the G0-G1 phase of the cell cycle compared to control cells in both the KLS and the MyPro compartments (Figure 5G-H). Complementarily, a smaller fraction of KLS and MyPro cells from CXCL12+ mice were in the synthesis (S) phase of the cell cycle compared to control cells (Figure 5G-H). Together, these results demonstrate that overexpression of *Cxcl12* in mice confers radiation protection to HSPCs by increasing quiescence of BM progenitor cells.

## DISCUSSION

In this study we generated and characterized a novel doxycycline-inducible *Cxcl12* transgenic mouse, and used this new tool to extensively assess the functional consequences of ubiquitous *Cxcl12* overexpression on hematopoietic function. Since the initial efforts to clone and determine the function of secreted cytokines and chemokines over two decades ago, various studies have highlighted the roles of CXCL12. These roles range from B-cell lymphopoiesis, hematopoietic cell survival and septal defects to HIV and cancer (Cavallero et al., 2015; Domanska et al., 2013; Guyon, 2014; Janssens et al., 2018; Meng et al., 2018; Nagasawa, 2007; Page et al., 2018; Patrussi and Baldari, 2011; Reid et al., 2018). Specifically, in hematopoiesis, CXCL12 acts as a chemoattractant for multiple cells, including B-cells and HSPCs, via its receptor CXCR4, which in itself is expressed to varying degrees by these cell types (Cordeiro Gomes et al., 2016; Ratajczak et al., 2006; Smith-Berdan et al., 2011). Here, we showed that altering *Cxcl12* expression does not affect the levels of its more predominant receptor *Cxcr4* or its novel receptor *Cxcr7* levels (Figure 1B). Additionally, we demonstrated that the migration of HSPCs towards the BM supernatant from CXCL12+ mice is increased, confirming higher amounts of functional secreted CXCL12 in the BM supernatant (Figure 1D). In adult mice, widespread overexpression of *Cxcl12* increased MPP numbers in the BM and spleen (Figure 2). While it is known that the conditional deletion of *Cxcl12* from IL-7-producing cells, a subset of leptin receptor-expressing cells, reduces both HSC and MPP numbers in the BM (Cordeiro Gomes et al., 2016), here we found that increased levels of CXCL12 led to a significant increase only in MPPs, but not HSCs, in both the BM and the spleen.

CXCL12 is produced by a variety of cells, including endothelial cells, perivascular cells and osteoblasts (Asada et al., 2017; Ding and Morrison, 2013; Ding et al., 2012; González et al., 2010; Mendelson and Frenette, 2014; Nagasawa et al., 1996; Pitt et al., 2015; Sugiyama et al., 2006). There is also evidence for sequestration of HSCs to the CXCL12-rich niche in the BM (Perlin et al., 2017; Reid et al., 2018; Sugiyama et al., 2006). Here, we observed that widespread overexpression of *Cxcl12* hindered long-term engraftment and reconstitution of the various mature hematopoietic cells (Figures 4C-D and S3C). Additionally, we infer that cells in the KLS compartment, i.e. cells that are high in the hematopoietic hierarchy, are more influenced by the CXCL12 overexpressing niche compared to more differentiated MyPros, during transplantation (Figure S3C versus S3D). Interestingly, the donor cells that have been pre-exposed to elevated CXCL12 levels and themselves have the capability to make CXCL12, also perform poorly when transplanted into a WT recipient (Figure 3B). It is notable that while the differences in the donor engraftment eventually disappear in the CXCL12+ hosts, they persist in the CXCL12+ donors compared to the controls (Figure 4D versus Figure 3B). This might imply that the CXCL12+ donor cells are possibly not reaching the supportive niche space for efficient LTMR due to the “microbubble CXCL12 overexpression” around them at the time of transplantation and hence do not engraft as well as the control cells in the WT environment.

Previous reports showed that the CXCR4-CXCL12 interaction is a key factor in the retention of the HSPCs in the BM. Loss of CXCL12 in the BM or inhibition of the CXCR4-CXCL12 interaction using AMD3100 led to both significantly reduced retention of HSPCs in the BM and facilitated their egress to the blood (Bonig et al., 2009; Dar et al., 2011; Redpath et al., 2017; Rosenkilde et al., 2004; Winkler et al., 2012). In our model where CXCL12 is broadly overexpressed, including in the blood (Figure 1Bii), BM specific CXCR4-CXCL12 interaction is perturbed, likely leading to the increased mobilization (Figures 5B, and S4C).

It has been reported that survival of myeloid progenitors in growth factor-deprived culture conditions is influenced by CXCL12 together with other niche factors, like stem cell factor (Broxmeyer et al., 2003b, 2003a). Additionally, previous work by various groups clearly demonstrated both that fewer cells are retained in the quiescent state in conditional *Cxcl12* knockout mice versus controls, and that loss or depletion of CXCL12 results in reduced quiescence of HSCs and altered hematopoietic regeneration upon 5-FU myelosuppression (Asada et al., 2017; Golan et al., 2013; Itkin and Lapidot, 2011; Tzeng et al., 2011). Our data obtained in the novel *Cxcl12* overexpression system support these results, whereby we distinctly observe higher numbers of cells in the G0 phase of the cell cycle and lower numbers in S phase (Figures 5G and H). Together, these findings suggest that the increased numbers of HSPCs in CXCL12+ mice is unlikely the result of increased cell proliferation, but more likely due to delayed differentiation. These slower cycling cells potentially also compete for niche space during transplantation and thus affect the LTMR potential of the donor cells in the CXCL12+ hosts (Figures 4D and S3C).

In addition to observing higher numbers of quiescent HSPCs, we also observed a radiation protective effect induced by *Cxcl12* overexpression (Figure 5D). Radiation protection can at least in part be explained by the predominance of cells in the quiescent state. Thus, the elevation of CXCL12 upon stress might be a self-preservation mechanism that cells adopt in emergency conditions. Radioprotection by CXCL12 might also be relevant in various cancers, in addition to the established role of CXCL12 in supporting epithelial-to-mesenchymal transition that results in cancer metastasis (Cheng et al., 2018; Cordeiro Gomes et al., 2016; Domanska et al., 2013; Meng et al., 2018; Ratajczak et al., 2006; Shan et al., 2015; Tang et al., 2019; Teicher and Fricker, 2010; Yu et al., 2017; Zhou et al., 2019). Using the inducible mouse model to test this will provide a better understanding of the consequence of elevated CXCL12 in a disease state and enable physicians to create better treatment plans.

Given the critical roles of CXCL12 in the control of HSPC localization, this novel mouse strain is a valuable tool to study hematopoietic trafficking *in vivo*. Notably, the ability to regulate *Cxcl12* overexpression in an inducible manner enables exquisite temporal control by modulating CXCL12 overexpression during various stages of development, either in early adulthood as performed here, or during early developmental stages or in aging. The extent of modulation and the effects in the latter scenario can facilitate understanding the critical role that *Cxcl12* plays in migration and seeding the different hematopoietic compartments with HSCs. The use of this novel tool could also shed light on the mechanisms underlying the roles of CXCL12 in hematopoietic development, including B-cell lymphopoiesis in both the fetal liver and BM and myelopoiesis (Nagasawa et al., 1996). In addition to temporal control, the CXCL12+ transgenic enables spatial or cell type-specific control of the CXCL12 expression. In this study we broadly expressed *Cxcl12* by crossing the *Cxcl12* transgenic mouse to the *Rosa rtTA* line. To obtain a controlled overexpression of *Cxcl12* in a cell type-specific manner, our new transgenic mouse can be crossed to a cell/tissue-specific *rtTA* strain or a *Lox-Stop-Lox-rtTA* mouse combined with a cell-specific *Cre* model. Since constitutive *Cxcl12* loss-of-function causes embryonic lethality, our new model provides a unique platform to conditionally re-express *Cxcl12* in a time- and tissue-specific manner on a null background during embryonic development (and later on), to study the diverse roles of *Cxcl12*.

## CONCLUSIONS

The novel *Cxcl12* overexpression mouse line that we have generated here exhibits salient features in hematopoiesis: reduced long-term multilineage reconstitution, amplified mobilization, and augmented radiation protection, which in turn is mediated by increased cell quiescence. In addition to its use in studies of hematopoiesis, this transgenic mouse strain can be utilized to understand the role of CXCL12 in various biological processes and tissues, including organ development, neural development and neurodegeneration, cardiovascular development and disease, inflammation, and cancer progression.

## MATERIALS AND METHODS

### Mice

Cxcl12 transgenic mice were generated at Cyagen (Santa Clara, CA) by pronuclear injection of the construct described in Figure S1A. Wild type (WT) Rosa rtTA (Cat# 006965) (Hochedlinger et al., 2005), WT C57BL6 (Cat# 000664), WT mT/mG (Cat# 007576) (Muzumdar et al., 2007) reporter mice, and WT UBC-GFP (Cat# 004353) (Schaefer et al., 2001) mice were purchased from Jackson Laboratories. Adult mice between 2-6 months of age and of both genders were used for the various experiments. All mice strains were maintained and bred at the UCSC animal facility and procedures were conducted according to the approved IACUC protocols. To genotype the *Cxcl12* transgenic mice, DNA was extracted and end-point PCR amplified for the transgenes using primers in Table S1.

Mice were provided doxycycline water prior to collection of a drop of blood from their tail and stained for the polymorphonuclear blood cells. Mice with both the *Cxcl12* and *rtTA* transgenes and expressing tomato in at least 10-15% of their cells were deemed CXCL12 overexpressing mice (CXCL12+).

### Antibodies, chemicals and other reagents

Doxycycline (Fisher Scientific) was resuspended at 1mg/mL in 5% sucrose (Fisher Technologies) water. Both control and CXCL12 transgenic mice were allowed to drink this doxycycline water ad libitum. Recombinant mouse CXCL12 was purchased from Peprotech. AMD3100 was purchased from Millipore Sigma. Antibodies (fluorophore conjugated, biotinylated, purified unconjugated, and fluorophore conjugated secondary antibodies including streptavidin conjugates) used for flow cytometry were purchased from BioLegend, BD Biosciences, eBioscience, and Life Technologies. CD117 magnetic beads (cKit beads for enrichment) and the MACS magnetic columns were purchased from Miltenyi Biotech.

### Real-time PCR

Messenger RNA was extracted from the various tissues using Trizol (Invitrogen) and quantified using a NanoDrop Spectrophotometer (ND-8000). Equal quantities of RNA were used to obtain cDNA using High Capacity cDNA Reverse Transcriptase Kit (Applied Biosystems) according to the manufacturer’s protocol. Quantitative real-time PCR was run on a ViiA 7 or QuantStudio 6 Flex PCR thermal cycler (Thermo Fisher Scientific) using SensiMix™ SYBR® No-ROX Kit (Bioline) according to the manufacturer’s protocol. β-actin was used to normalize transcript levels. Primers used for the real time PCR are in Table S2.

### Transwell *in vitro* migration assay

In vitro migration assays were performed as described before (Smith-Berdan et al., 2011). Briefly, cell-free BM supernatant was obtained from the control and CXCL12 transgenic mice by crushing the long bones and centrifuging the sample. BM cells from non-doxycycline treated mice were isolated and lineage depleted using Sheep Anti-Rat IgG magnetic Dynabeads (Invitrogen). The lineage depleted cells were heat activated for an hour at 37°C prior to placing them on the 5μm pore transwell insert. The cell-free BM supernatant was used as the chemoattractant in the lower chamber of the 24-well plate. Migration of the cells took place at 37°C for 2 hours in an incubator with 5% CO_2_. Post migration, APC Calibrite beads (BD Biosciences) were added to the lower chamber and cells were collected, stained and analyzed by flow cytometry.

### Flow Cytometry

To perform flow cytometry, the cells were isolated and stained with respective antibodies in staining media (5mM EDTA in 1X PBS with 2% heat inactivated donor calf serum) on ice and in the dark. The washed and pelleted cells were then resuspended in staining media with propidium iodide to exclude dead cells. Cells were filtered through a 70μm nylon mesh and run on a FACSAriaIII flow cytometer (BD Biosystems) and analyzed using FlowJo software (Beaudin et al., 2016; Boyer et al., 2019; Smith-Berdan et al., 2015). The phenotypic markers used for identification of the different cell types are as follows:

cKit+: Lineage- cKit+;

HSC: Hematopoietic stem cells (Lineage- cKit+ Sca1+ Flk2- CD150+);

MPP: Multipotent progenitor cells (Lineage- cKit+ Sca1+ Flk2+ CD150-);

MyPros: Myeloid progenitor cells (Lineage- cKit+ Sca1-);

CMP: Common myeloid progenitors (Lineage- cKit+ Sca1- FcγRα_mid_ CD34_mid_);

GMP: Granulocyte-Macrophage progenitors (Lineage- cKit+ Sca1- FcγRα_high_ CD34_high_);

MEP: Megakaryocytic-Erythroid progenitors (Lineage- cKit+ Sca1- FcγRα_low_ CD34_low_);

GM: Granulocyte-Macrophage cells (Ter119- Mac1+ Gr1+ B220- CD3-);

CLP: Common Lymphoid progenitors (Lineage- IL7Rα+ Flk2+ cKit_mid_ Sca1_mid_);

B: B-cells (Ter119- Mac1- Gr1- B220+ CD3-);

T: T-cells (Ter119- Mac1- Gr1- B220- CD3+);

MSC: Mesenchymal stem cells (CD45- Ter119- CD31- Sca1+ CD51+);

OBL: Osteoblast cells (CD45- Ter119- CD31- Sca1+ CD51-);

EC: Endothelial cells (CD45- Ter119- CD31+ Sca1+);

CD27 marker was used to define HSCs and MPPs in the spleen and blood tissues.

### Proximity Ligation Assay (PLA) for detection of individual molecules of CXCL12

Control and *Cxcl12* transgenic mice were sacrificed and the femur and tibia were placed in 4% methanol-free PFA (Thermo Fisher) for 21 hours. They were washed in 1X PBS (without Ca_++_, Mg_++_) for 1 hour. This was repeated once more and the hindlimb bones were then placed in fresh PBS. Bones were then decalcified in 10% EDTA, pH 8 for 14 days. Tissues were embedded in 4% low temperature gelling agarose and cut into 100-150 micrometer thick section using a vibratome (Leica VT1200 S). The Proximity Ligation Assay (PLA) was carried out as described in Kunz et al (Kunz and Schroeder, 2019). Briefly, after applying the primary anti-CXCL12 (ab9797, Abcam) and the anti-CD31 (AF3628; R&D Systems) antibodies overnight at room temperature, the “Far-red” detection kit was used to visualize CXCL12 bound antibody by the PLA. Unspecific binding of fluorescently-labeled oligonucleotides in BM was blocked by addition of the oligonucleotide ODN 1826 during blocking and permeabilization of the tissue sections. One section per mouse was imaged and 4 ROIs (1024 by 1024 pixels at 63x magnification) per section were quantified. The number of CXCL12 signal i.e. the number of spots within the tissue volume, given by DAPI, was determined and normalized to the tissue volume analyzed.

### Tissue Extraction

Blood from the entire mouse was obtained by perfusing the mouse with 20mM EDTA in 1X PBS. The BM cells were obtained by crushing the tibia and femur (and occasionally, hip bones). Bone stromal cells were obtained by treating the crushed bone with collagenase type 1 (Worthington) enzyme (3mg/mL) at 37°C for an hour with intermittent vortexing (Smith-Berdan et al., 2012). Splenic cells were obtained by crushing the spleen in staining media. Lung tissue was digested with collagenase type 1 (Worthington) enzyme (800U/mouse) for 45 minutes at 37°C with intermittent vortexing. Liver was digested with collagenase type IV (Worthington) enzyme (1mg/mL) and DNAse I (Millipore Sigma) enzyme (100U/mL) (Leung et al., 2019).

### Transplantation assays

For long-term engraftment and multilineage reconstitution assays, either whole bone marrow (WBM) cells or cKit enriched double sorted cells from the UBC-GFP or the control/CXCL12 transgenic mice were transplanted retro-orbitally into sub-lethally (525 rads) irradiated (Faxitron CP-160, Precision Instruments) recipients. Peripheral blood was obtained at various times post-transplantation, for up to 4 months, and stained and analyzed for the different mature cell populations as described before (Boyer et al., 2019). Short-term homing assays were performed by retro-orbital injection of the KLS cells obtained from cKit enriched, double sorted (FACSAriaIII) GFP+ BM cells, into lethally irradiated (2 doses of 525 rads, with a three-hour separation between the doses) recipients. Three hours post-transplantation, different tissues were extracted and the GFP+ donor cells were assessed by flow cytometry (Boyer et al., 2011; Smith-Berdan et al., 2011, 2015).

### Vascular permeability assay

Miles assay with Evans Blue was performed as described (Miles et al., 1952; Smith-Berdan et al., 2019). In short, Evans Blue (50mg/kg) was injected intravenously into the mice, which were euthanized after 5 minutes. The weights of the different tissues were calculated and then the tissues were placed overnight in formamide (Sigma) at 55°C to allow the release of the dye. The absorbance of the dye (in the formamide) at 650nm per gram of the tissue provided a measure of the vascular permeability.

### Spleen CFU assay

One hundred double sorted GFP+ HSCs from the BM were transplanted retro-orbitally into lethally irradiated (2X of 525 rads with a three-hour gap) control/CXCL12 transgenic mice that were maintained on dox at least 7 days prior to transplantation. The spleens were removed from the recipients 12.5 days post-transplantation and examined (Boyer et al., 2011, 2019; Smith-Berdan et al., 2011). The number of individual donor-derived colonies were counted and dissected under a fluorescence microscope. Single cell suspensions of the colonies were stained and analyzed on a cytometer to evaluate the different cell types (Boyer et al., 2019):

EP (CFU-S): Erythroid progenitors (FSC_mid-hi_ Ter119+ CD41- Mac1- Gr1- B220-);

Meg (CFU-S): Megakaryocyte progenitors (FSC_mid-hi_ Ter119- CD41+ Mac1- Gr1- B220-);

GM (CFU-S): Granulocyte-Macrophage (FSC_mid-hi_ Ter119- CD41- Mac1+ Gr1+ B220-);

B (CFU-S): B-cells (FSC_mid-hi_ Ter119- CD41- Mac1- Gr1- B220+).

### Mobilization assay

Mice were injected subcutaneously once with AMD3100 (5mg/kg) as described (Smith-Berdan et al., 2019). One-hour post injection, mice were euthanized and perfused to obtain blood. After red blood cell lysis, cells were stained and analyzed by flow cytometry.

### Radiation sensitization assay

Both control and CXCL12 transgenic mice were irradiated (525 rads) using an X-ray tube irradiator (Faxitron CP-160) after being on doxycycline for at least 7 days. 7- or 11-days post irradiation (mice continued to get doxycycline), the mice were perfused, the different tissues were extracted and the cells analyzed by flow cytometry.

### Cell cycle analysis

Cell cycle analysis was performed using the Click-iT™ Plus EdU Alexa Fluor™ 647 Flow Cytometry Assay Kit (Thermo Fisher Scientific) according to the manufacturer’s protocol (Beaudin et al., 2014, 2016; Ugarte et al., 2015). Briefly, mice were irradiated (525 rads), and EdU was injected 7 days after irradiation. Mice were euthanized 30 minutes after EdU injection and perfused. BM cells were then incubated with cKit beads to enrich for HSPCs, stained for the different stem and progenitors’ cells, fixed, permeabilized, treated with azide for EdU incorporation and stained with Hoechst to obtain cell cycle status.

### Statistical analysis

All mice were maintained on doxycycline for at least 7 days prior to tissue extraction, unless specified otherwise.

Statistical significance was determined by unpaired two-tailed Student’s T-test when comparing two groups and by one-way ANOVA followed by post-hoc Tukey test when comparing more than two groups unless stated otherwise (Boyer et al., 2019; Rajendiran et al., 2016; Smith-Berdan et al., 2019). All the data are represented as mean ± s.e.m. The sample size, number of independent experiments, and P-values are all provided for each experiment in the respective figure legend.

## ACKNOWLEDGEMENTS

We thank the Brennen Cooper and Jackson Hauty for genotyping assistance. We thank the University of California, Santa Cruz Vivarium Staff and Bari Nazario from the Institute for the Biology of Stem Cells (IBSC) Flow Core and Cell Culture Facilities.

## Author contributions

S.R. and C.E.F. conceived of the study, designed experiments and wrote the paper. S.R., S.S.B., L.K., M.R., L.S. and T.S. performed experiments and analyzed and interpreted data. All authors read and approved the final manuscript.

